## Supplemental Figures and Tables for "Substrate inhibition imposes fitness penalty at high protein stability"

Supplementary Figures and Table:

4 Figures and 1 Table

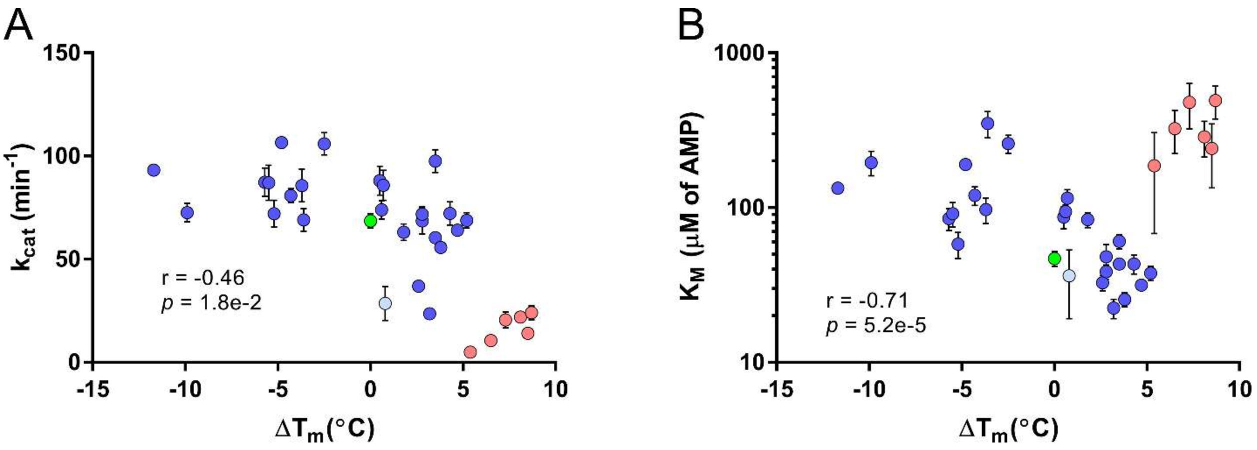


**Fig S1: Activity parameters vs stability.** Correlation between $k_{cat}$ (A) or $K_M$ (B) and protein stability is shown. Log values of $K_M$ were used for correlation calculations. The WT is shown in green, L82V in light blue, whereas all mutants involving mutation at Q16 position are shown in pink circles. The error bars are s.e.m. of three measurements. In both panels, Pearson correlations are calculated without considering Q16 mutants (red points).

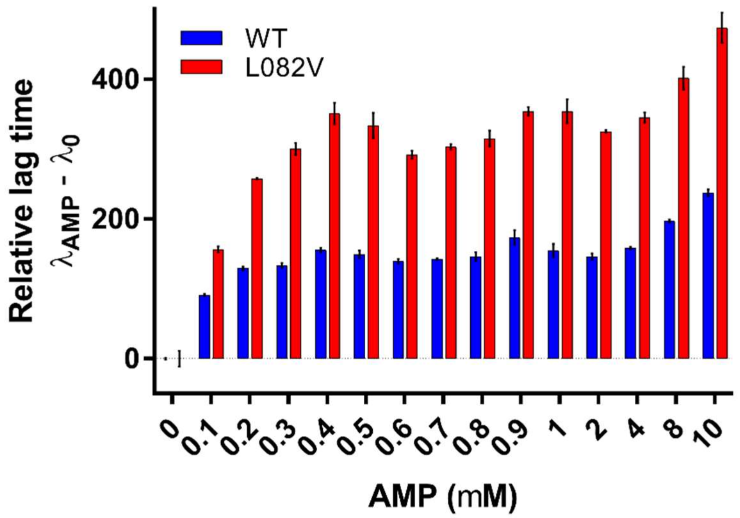


**Fig S2: Pilot experiment to estimate the dynamic range of AMP effect on fitness.** A pilot experiment with only WT and L82V, the most inhibited mutant in this study, were grown in M9 media containing various amounts of AMP. Relative lag times increase substantially up to 400 μM of AMP, following which the changes are smaller. The error bars are s.e.m. from three colonies.

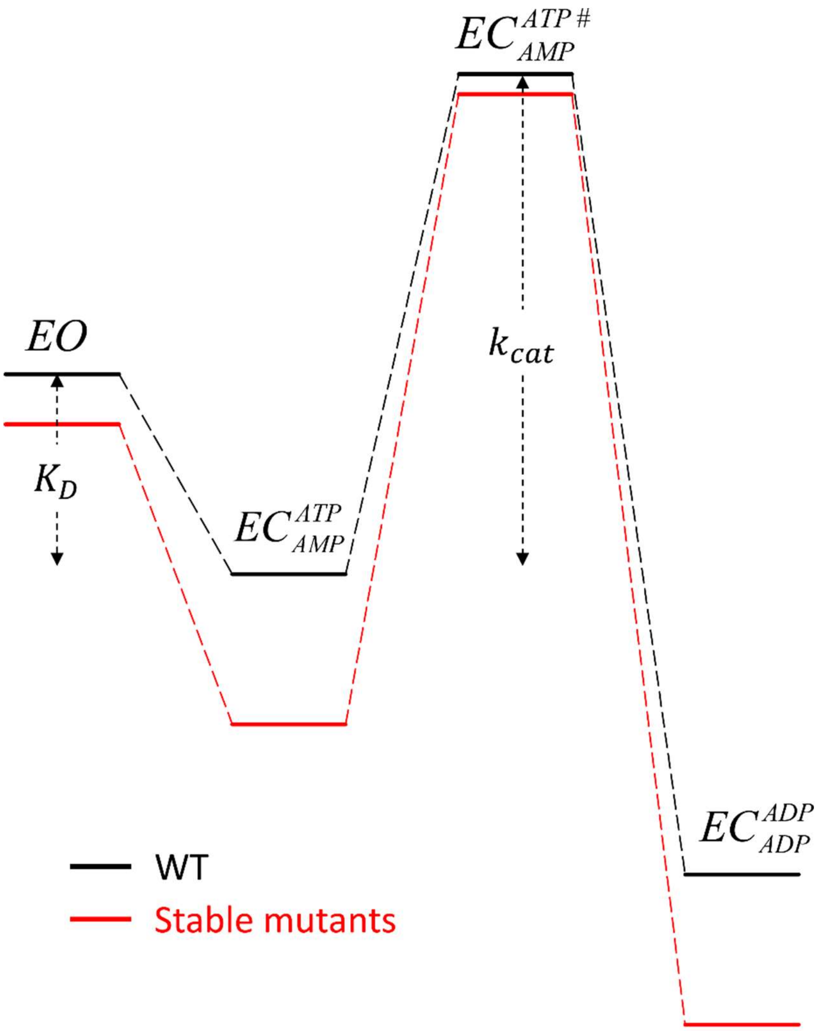


**Fig S3: Schematic of possible effects of stabilization on enzyme mechanics.** The schema depicts an energy landscape during catalysis. WT scheme is shown in black lines, whereas that of a stabilizing version is in red. EO and EC are the 'open' and 'closed' states of the enzyme. The more stable enzyme may preferentially stabilize the ligand-bound closed state more than the unbound open state, which will be reflected in stronger $K_D$ and hence stronger $K_M$. Such preference may also result in a high activation barrier $\left(EC_{AMP}^{ATP} \rightarrow EC_{AMP}^{ATP\#}\right)$, which results in reduced $k_{cat}$.

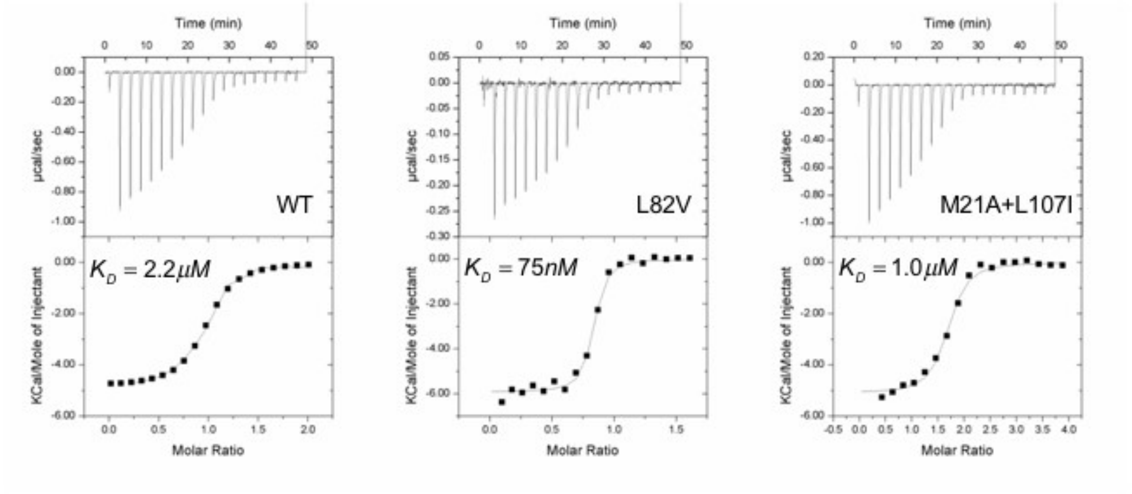


**Fig S4: Ap5A binding by ITC.** Binding of a bidentate inhibitor Ap5A was measured by ITC to WT, L82V, and M21A+L107I. The concentration of protein used in the cell was 96.7 µM of WT, and 24.1 µM each of the two mutants. 1 mM of the inhibitor was used in syringe for WT and M21A+L107I, whereas 200 µM was used for L82V titrations. The binding measurement was done at 25 °C.

**Table S1: Growth parameters of Adk strains**

| Mutant | Growth rate ($min^{-1}$) | | Lag time (min) | |
|---|---|---|---|---|
| | mean | SEM | mean | SEM |
| V106W | 0.0087 | 0.0001 | 240.4 | 6.1 |
| Q016F | 0.0107 | 0.0001 | 248.0 | 1.0 |
| Q016F + V169E | 0.0099 | 0.0001 | 259.3 | 1.3 |
| V106H | 0.0126 | 0.0001 | 189.7 | 2.4 |
| L083I | 0.0091 | 0.0001 | 240.9 | 4.2 |
| A093L | 0.0091 | 0.0001 | 218.4 | 10.2 |
| A093I | 0.0093 | 0.0002 | 243.3 | 2.4 |
| L082F | 0.0132 | 0.0001 | 233.2 | 3.5 |
| WT | 0.0133 | 0.0003 | 239.1 | 4.1 |
| M021A | 0.0130 | 0.0004 | 241.2 | 4.3 |
| L083F | 0.0112 | 0.0001 | 235.9 | 2.8 |
| V169E | 0.0130 | 0.0009 | 241.0 | 3.5 |
| L107I | 0.0121 | 0.0007 | 248.9 | 4.1 |
| M021A + V169E | 0.0143 | 0.0004 | 248.7 | 2.2 |
| L107I + V169E | 0.0137 | 0.0003 | 234.8 | 1.9 |
| L209I | 0.0102 | 0.0002 | 246.3 | 7.3 |
| M021A + V169E + L209I | 0.0099 | 0.0003 | 235.8 | 6.5 |
| L082V | 0.0100 | 0.0001 | 268.7 | 6.0 |

All experiments were done in triplicates with at least three colonies (biological repeats)
